# Forward genetics reveals microsporidian drug targets and recombination-based resistance

**DOI:** 10.64898/2026.01.30.702766

**Authors:** Qingyuan Huang, Xianzhi Meng, Winnie Zhao, Guoqing Pan, Jie Chen, Aaron W. Reinke

## Abstract

Microsporidia are a diverse group of intracellular fungal parasites that infect humans and many agriculturally important animals. Several microsporidia inhibitors have been characterized, but the targets of these inhibitors in microsporidia have not been confirmed, and the mechanisms by which microsporidia evolve resistance are unknown. Using the model organism *Caenorhabditis elegans* and its naturally occurring microsporidian parasite *Nematocida parisii*, we developed a technique to continually passage infected animals for multiple generations under increasing inhibitor concentrations. We obtained multiple independent *N. parisii* strains that are specifically resistant to the microsporidia inhibitors albendazole or dexrazoxane. The albendazole-resistant strains all contained either heterozygous or homozygous mutations in beta-tubulin, with several strains containing beta-tubulin variants that have been observed in other albendazole-resistant organisms. Strains containing homozygous variants are resistant to several albendazole analogs, whereas the heterozygous variant-containing strains are only resistant to albendazole. Several of the albendazole-resistant strains also contain mutations in alpha-tubulin. The dexrazoxane-resistant strains all contain heterozygous mutations in topoisomerase II and several of these mutations occur in the binding site of this inhibitor. By mapping heterozygosity of the resistant strains across the *N. parisii* genome, we observed loss of heterozygosity spanning the beta-tubulin locus in the albendazole-resistant strains containing homozygous beta-tubulin variants. Our study demonstrates the utility of forward genetics for identifying microsporidian drug targets. We also find that drug resistance arises through *de novo* heterozygous mutations that can subsequently become homozygous, likely via mitotic recombination.

## Introduction

Microsporidia infect humans and agriculturally important animals such as honeybees, silkworms, and shrimp [1,2]. There are at least 17 species known to infect humans and cause symptoms such as diarrhea, including reports of lethal infections [3]. Microsporidia infections in silkworms have impacted the silk industry since the 1850s and more recently infections in shrimp have been reported to cause hundreds of millions of dollars in losses each year [4,5]. Because of the threat of microsporidia to health and food security, there is a need for effective microsporidia inhibitors. The two most commonly used microsporidian drugs are fumagillin and albendazole, although host toxicity has limited the use of fumagillin and several species are resistant to albendazole [6–9].

The model organism nematode *Caenorhabditis elegans* and its naturally occurring microsporidian parasite *Nematocida parisii* have been developed as a convenient model for identifying microsporidia inhibitors and determining their mechanisms [10–12]. Infection of *C. elegans* begins when a microsporidian spore is ingested and the unique infectious apparatus known as the polar tube is used to deposit a sporoplasm inside an intestinal cell [13]. This sporoplasm then divides into an intracellular stage known as a meront, the meronts proliferate in intestinal cells, and by three days have differentiated into spores [14]. Infection by *N. parisii* prevents *C. elegans* progeny production, providing an easy-to-screen phenotype [12]. Using 96-well plates, over 10,000 compounds have been tested, with the identification of inhibitors that inactivate spores, block spore germination, and prevent proliferation [12,15–17]. Although the stage at which these inhibitors act has been elucidated, the molecular target of any of these inhibitors has not been determined.

How microsporidia evolve resistance to drugs is unknown. Although fumagillin has been administered for decades to control microsporidia infections in honey bees, there is no evidence that resistance has occurred [18]. Albendazole, which belongs to a class of inhibitors called benzimidazoles, is thought to inhibit beta-tubulin and several human-infecting species contain variants hypothesized to provide benzimidazole resistance [6,7]. Dexrazoxane is a member of a class of inhibitors called bisdioxopiperazines [19]. This drug is used to prevent the toxic effects of anthracycline chemotherapy, and is thought to function both through its ability to chelate iron and act as a catalytic inhibitor of topoisomerase II [20]. We previously demonstrated that dexrazoxane is unlikely to inhibit *N. parisii* proliferation through iron chelation [12].

To determine the molecular target of microsporidia inhibitors, we have developed an approach to isolate and characterize drug-resistant strains of *N. parisii*. Using large-scale infections of *C. elegans* with *N. parisii*, we treated cultures with increasing amounts of the drugs albendazole and dexrazoxane over multiple generations to select for drug-resistant strains. All the resistant strains are specific for the drug they were selected for, with some albendazole-resistant strains also being resistant to other benzimidazoles. All albendazole-resistant strains contain heterozygous or homozygous variants in beta-tubulin, with two of these strains also containing variants in alpha-tubulin. Several of these beta-tubulin variants are in positions that have been observed in albendazole-resistance in other organisms, including a variant in a position that has been implicated in providing resistance in several human-infecting microsporidia species. The dexrazoxane-resistant strains all contain heterozygous variants in topoisomerase II, and several of these variants occur in the dexrazoxane binding site. Heterozygosity is lost in the homozygous beta-tubulin variant containing strains on the chromosome encoding beta-tubulin. Together our results demonstrate the ability to determine drug targets in microsporidia and show that evolution of resistance to inhibitors is through the spontaneous generation of heterozygous variants, as well as through homozygous variants which likely occur through mitotic recombination.

## Results

### Isolation of albendazole-resistant *N. parisii* isolates

The isolation of rare mutations requires the ability to culture large numbers of an organism. We have previously established small-scale liquid infections for screening compounds with inhibitory ability towards *N. parisii* [12,15–17]. We first tested using an intermediate volume, 3 mL, of a media used for long-term culture of *C. elegans* (Fig. 1A). After infection for four days at 21°C in the presence of various albendazole concentrations, we fixed worms and stained with the dye direct yellow 96 (DY96), which binds to the chitin-containing microsporidia spores and *C. elegans* embryos (Fig. 1B). We quantified the percentage of animals with moderate-to-severe infection, which we defined as at least half of the intestine displaying newly generated spores. We observed no inhibition at 2.5 μM albendazole, intermediate levels of inhibition at 5 μM and 10 μM, and strong inhibition by 20 μM (Fig. 1C). These experiments demonstrate that infection can be monitored in larger volumes and identify doses of albendazole to be used to select resistant strains.

**Figure 1.**
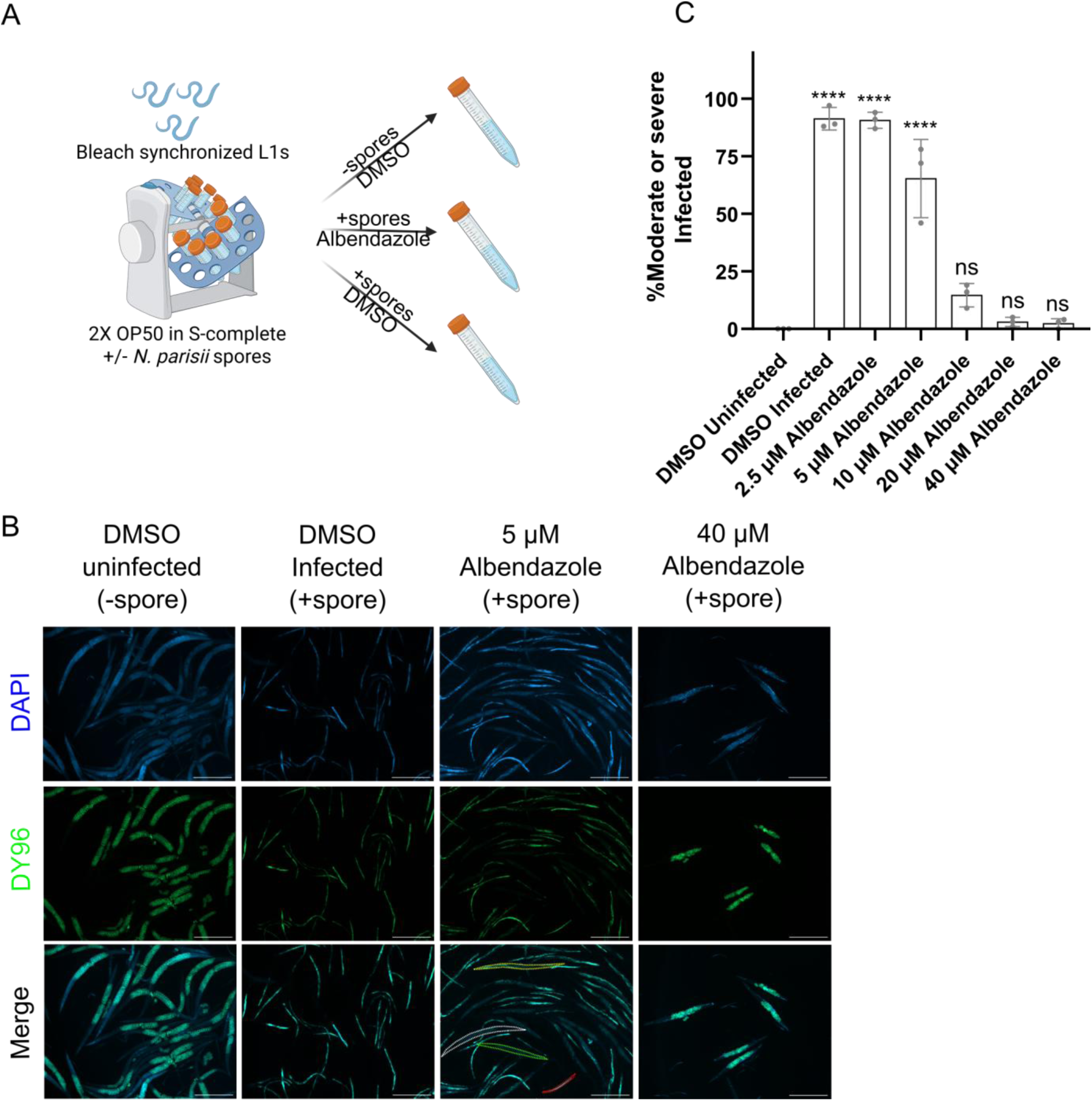
Establishment of liquid culture of *N. parisii* to obtain drug-resistant isolates. **A.** Schematic of the procedure for culturing *N. parisii* spores in liquid medium. 6,000 bleach-synchronized L1 animals were incubated in 3 mL of S-complete media with or without different concentrations of albendazole for 4 days at 21°C in the presence of 10,000 *N. parisii* spores/µL. **B.** Representative images (5× magnification; scale bar, 500 μm). (Left) Worms developed normally in the absence of spores and are gravid with embryos that are stained green with DY96 dye. (Middle-left) Microsporidia infection resulted in new spore formation (stained green with DY96 dye) and impaired development, preventing worms from reaching the gravid adult stage. (Middle-right & Right) Treatment with albendazole reduced new spore production and restored development to gravid adults. White dashed outlines: uninfected worms; green: lightly infected; yellow: moderately infected; red: severely infected. **C.** Percentage of animals showing moderate or severe infection (n = 3 biological replicates; N ≥ 100 animals counted per replicate). Significance was evaluated relative to DMSO-uninfected controls using a one-way ANOVA with Dunnett’s post-hoc test: **** p < 0.0001; ns, not significant (p > 0.05).

To select for rare resistant mutants, we scaled up this culture system. Using the conditions established above, we set up three independent flasks of worms infected with *N. parisii.* For each replicate, 10,000 bleach-synchronized L1 worms were cultured with *N. parisii* spores in 50 mL of media containing 5 μM albendazole. After four days, the population infection rate was quantified. The albendazole concentration was decreased if >80% of adults were uninfected, increased if >50% showed moderate-to-severe infection, and maintained otherwise (Fig. 2A). After 9 generations, none of the initial three flasks appeared to have resulted in emerging resistance (Fig. S1A, C, and E). We then used these same three flasks and increased the culturing time for each generation from four days to six days. To increase the *N. parisii* growth rate, we also increased the temperature to 25°C [13,21]. After 4-11 more generations, all three flasks displayed resistance to albendazole, which we defined as cultures exhibiting >50% moderate-to-severe infection at 60 μM albendazole (Fig. S1B, D, and F). We then set up an additional three flasks to select for albendazole resistance, using a 6-day incubation time at 25°C. All three flasks displayed the emergence of resistance within 10-16 generations (Fig. S1G-I).

**Figure 2.**
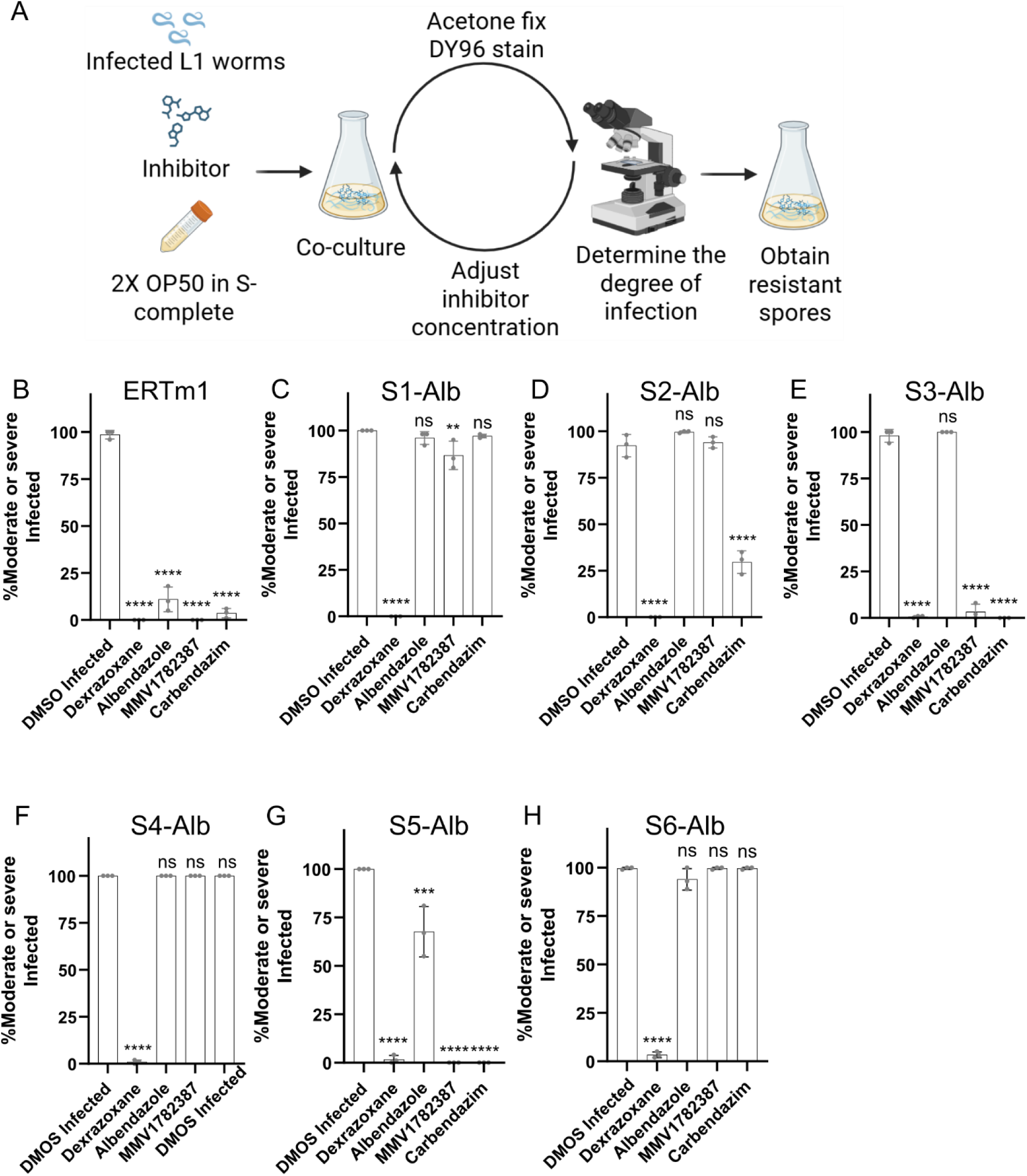
Isolation and characterization of albendazole-resistant strains of *N. parisii.* **A.** Workflow for culturing drug-resistant spores. Infected worms were co-cultured with bleach-synchronized L1 animals, and the inhibitor concentration was adjusted according to the severity of infection. **B–H**. Spore preparations were generated for drug-resistant strains of *N. parisii*, and these strains were tested using a 24-well continuous infection assay. The assay conditions include 10,000 spores/μL of *N. parisii*, 4 days of infection at 21°C, and 60 μM of the indicated benzimidazole and 120 μM dexrazoxane. Percentage of animals showing moderate to severe infection in ERTm1 and albendazole-resistant groups (n = 3 biological replicates; N ≥ 100 animals per replicate). Significance was evaluated relative to DMSO-infected controls using one-way ANOVA with Dunnett’s post-hoc test: **p < 0.01, ****p < 0.0001; ns, not significant (p > 0.05).

To confirm the resistance phenotype of selected *N. parisii* isolates, we generated spore preparations for each selected strain, and assayed for resistance using our standard 24-well infection assay [17]. We used an infection dose so that >90% of worms were infected with *N. parisii*. For the wild-type ERTm1 strain, addition of 60 μM albendazole resulted in ∼10-fold decrease in infection (Fig. 2B). For each of the six isolated albendazole-resistant strains, S1-Alb to S6-Alb, treated with 60 μM albendazole, all showed either no decrease in infection or a slight decrease in infection in the case of S5-Alb (Fig. 2C-H). To determine if this resistance was specific to albendazole, we tested the wild-type spores and the six resistant strains with the microsporidia inhibitor dexrazoxane. All strains showed complete inhibition with 120 μM dexrazoxane (Fig. 2B-H). Together these results demonstrate the successful isolation of *N. parisii* strains with specific resistance to albendazole.

To determine if the albendazole-resistant strains were specific to albendazole over other benzimidazoles, we tested two related analogs, carbendazim and MMV1782387, that were previously shown to have inhibitory activity towards *N. parisii* [15]. These three benzimidazole compounds differ in their benzimidazole-ring substituents: albendazole has a propylthio group, MMV1782387 has a single methyl group, and carbendazim has an unsubstituted benzimidazole ring. S1-Alb, S4-Alb, and S6-Alb displayed complete resistance to carbendazim (Fig. 2C, F, and H). S4-Alb and S6-Alb also displayed complete resistance to MMV1782387, whereas S1-Alb was resistant to MMV1782387, though not to the same extent as to the other two benzimidazoles (Fig. 2C, F, and H). In contrast, S3-Alb and S5-Alb were not resistant to MMV1782387 or carbendazim (Fig. 2E and G). S2-Alb was resistant to MMV1782387 and partially resistant to carbendazim (Fig. 2D). These results show that although all strains are resistant to albendazole, there are differences in specificity towards other benzimidazoles.

### Isolation and characterization of dexrazoxane-resistant *N. parisii* isolates

To determine if our approach to generate drug-resistant *N. parisii* spores could be used on other inhibitors, we applied our approach to dexrazoxane. We performed three independent selections for dexrazoxane resistance, using the modified conditions of a 6-day generation time and culturing temperature of 25°C. We used a starting concentration of 32 μM dexrazoxane and after 6-13 generations, all three independently derived strains showed over 50% of worms displaying moderate or severe infection at 120 μM dexrazoxane (Fig. S2).

We then characterized the dexrazoxane-resistant *N. parisii* strains. We generated spore preparations for the three resistant strains (S1-S3-Dex). Under conditions where greater than 80% of worms showed moderate or severe infection, 120 μM dexrazoxane prevented infection by the wild-type ERTm1 strain (Fig. 3A). Each of the three dexrazoxane-resistant strains did not display any significant inhibition with 120 μM dexrazoxane (Fig. 3B-D). To determine the specificity of these strains, we also tested them with albendazole and all the strains displayed wild-type levels of sensitivity to this inhibitor (Fig. 3A-D). Together these experiments demonstrate the successful isolation of *N. parisii* strains with specific resistance towards dexrazoxane.

**Figure 3.**
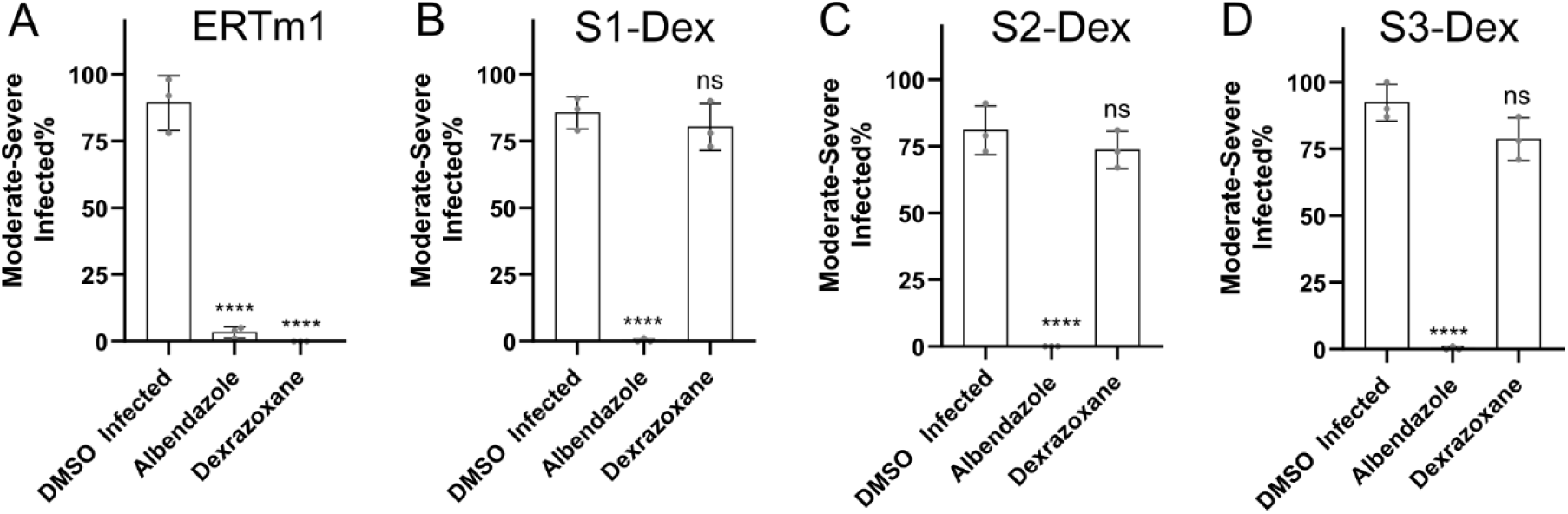
Isolation of dexrazoxane-resistant *N. parisii* strains. A–D. 24-well continuous infection assays were conducted under the following conditions: 10,000 spores/μL of *N. parisii*, a 4-day infection period at 21°C, and 60 μM albendazole or 120 μM dexrazoxane. Percentage of animals showing moderate to severe infection in the ERTm1 and dexrazoxane-resistant strains (n = 3 biological replicates; N ≥ 100 animals per replicate). Significance was evaluated relative to DMSO-infected controls using one-way ANOVA with Dunnett’s post-hoc test: **p < 0.01, ****p < 0.0001; ns, not significant (p > 0.05).

### Beta-tubulin mutations occur in all albendazole-resistant *N. parisii* isolates

To determine the putative causative mutations in our drug-resistant *N. parisii* strains, we performed whole genome sequencing of the six albendazole-resistant isolates, three dexrazoxane-resistant isolates, and two wild-type ERTm1 samples. To improve mapping of sequence reads from the resistant *N. parisii* strains, we generated a long-read-based genome. This genome assembly contained seven contigs. We then mapped reads from all of our samples to this genome assembly. We determined both homozygous and heterozygous variants of each strain, retaining only those with a predicted protein-coding impact. We then compared all the albendazole or dexrazoxane-resistant strains to the corresponding non-resistant strains to determine variants that were specifically associated with either albendazole or dexrazoxane resistance. Two of the strains, S4-Alb and S6-Alb, had identical variant patterns and do not represent independent occurrences. Because of this, we consider S4-Alb and S6-Alb as a single resistant strain. We detected between 0-2 homozygous variants and 1-15 heterozygous variants per strain (Table S1). On average, we detected ∼7 variants per strain and we did not observe any identical variants that occurred in multiple independently selected strains.

All albendazole-resistant strains contained non-synonymous variants in beta-tubulin. We observed homozygous mutations in strain S1-Alb (E198K), S2-Alb (F20L), and S4/S6-Alb (Q134K) (Fig. 4). We also observed heterozygous variants in S3-Alb (A248T and A314T) and S5-Alb (A314V) (Fig. 4). The albendazole-resistant strains also contained several variants in alpha-tubulin, with a homozygous variant in S2-Alb (Y85H), and a heterozygous variant in S5-Alb (E251D) (Fig. S4). The only other genes that contained variants in multiple strains were genes encoding either short or repetitive proteins, with only two strains containing a variant, and at least one of these variants was a deletion, frameshift, or insertion (Table S1). To validate the observed variants of beta-tubulin, we performed PCR and sequenced the beta-tubulin gene. We detected the corresponding homozygous variants in S1-Alb, S2-Alb, S4-Alb, and S6-Alb, as well as the heterozygous variants in S3-Alb and S5-Alb (Fig. S3).

**Figure 4.**
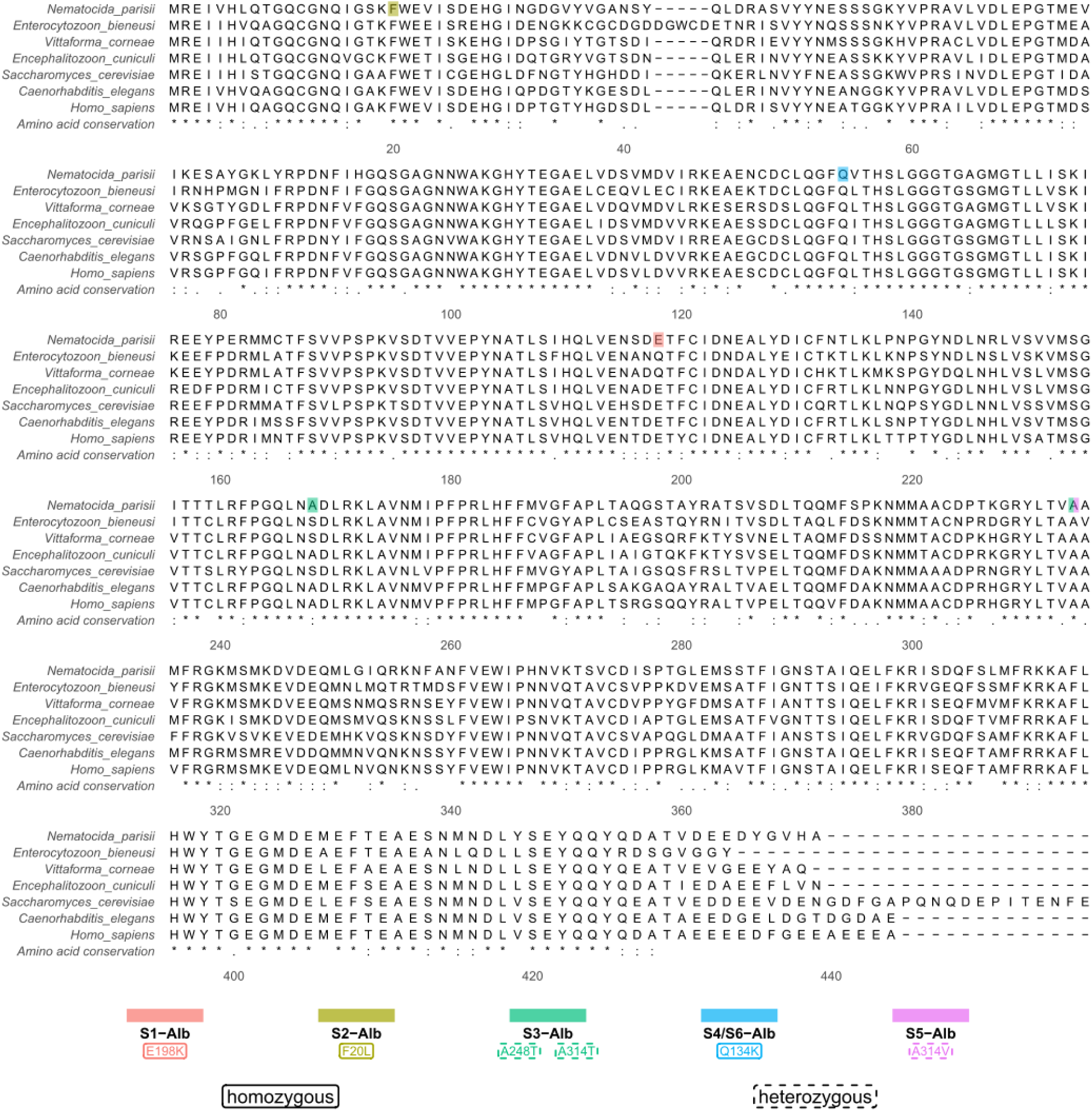
Albendazole-resistant *N. parisii* strains contain either homozygous or heterozygous variants in beta-tubulin. Multiple sequence alignment of beta-tubulin sequences from microsporidia *(N. parisii*, *E. bieneusi*, *V. corneae*, and *E. cuniculi), S. cerevisiae*, *C. elegans*, and humans. Homozygous and heterozygous variants detected in the five independently selected albendazole-resistant strains are annotated according to the legend at the bottom. Positions are numbered according to *S. cerevisiae* TUB2 and amino acid conservation is shown at the bottom of the alignment.

### Dexrazoxane-resistant *N. parisii* isolates contain heterozygous variants in topoisomerase II

We then analyzed variants that were specific to the dexrazoxane-resistant strains. We observed heterozygous variants in topoisomerase II for strains S1-Dex (N142S), S2-Dex (Y144H), and S3-Dex (V243M) (Fig. 5). Positions 142 and 144 are in the dexrazoxane binding site in the budding yeast ortholog of topoisomerase II, and variants in the corresponding position of 144 for the human ortholog provide dexrazoxane resistance (Fig. 5) [22,23]. No genes besides topoisomerase II contained variants in more than one strain (Table S1). We also detected a homozygous variant in S1-Dex in the gene elongation factor 2 (Table S1).

**Figure 5.**
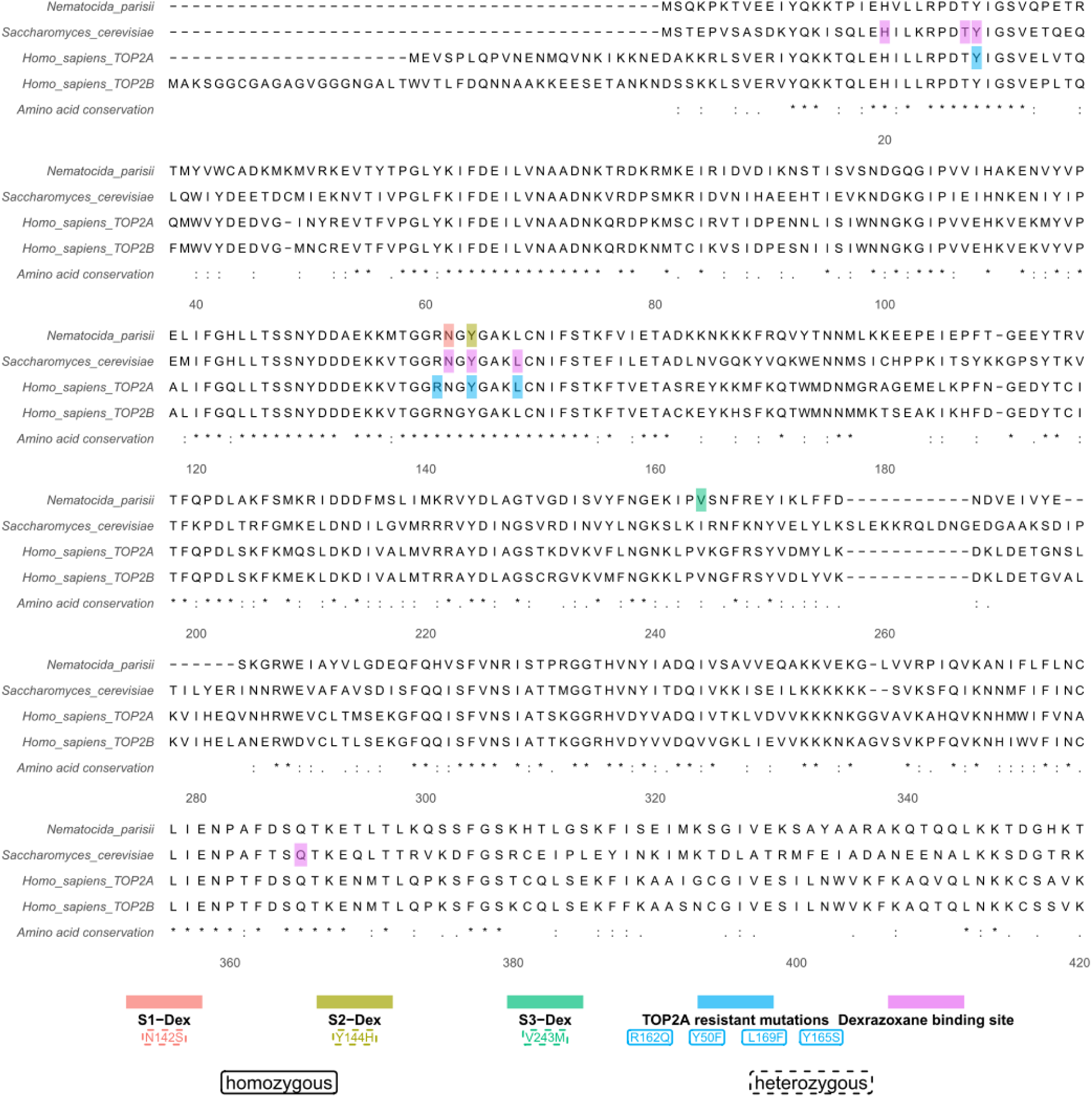
Dexrazoxane-resistant *N. parisii* strains contain heterozygous variants in topoisomerase II. Multiple sequence alignment of the ATPase domain of topoisomerase II (Residues 1-413 of *S. cerevisiae* TOP2 [22]) for *N. parisii*, *S. cerevisiae*, and humans. Heterozygous variants detected in the three dexrazoxane-resistant *N. parisii* strains are shown according to legend at the bottom. Mutations in human TOP2A that result in bisdioxopiperazine resistance and the amino acid position from the dexrazoxane binding pocket in *S. cerevisiae* TOP2 are annotated according to the legend at the bottom [22,23,43,62–64]. Positions are numbered according to *S. cerevisiae* TOP2 and amino acid conservation is shown at the bottom of the alignment.

### Loss of heterozygosity in some *N. parisii* albendazole-resistant isolates spanning the beta-tubulin locus

The observation of multiple independent homozygous mutations in beta-tubulin suggests the occurrence of some large-scale genomic alterations. To determine if there were copy number differences of chromosomes, we mapped the read depth across the *N. parisii* assembly. We observed similar read depth in both our wild-type and drug-resistant *N. parisii* strains across the seven contigs, providing evidence that aneuploidy does not explain the homozygous alleles observed (Fig. S5). An alternative explanation would be that recombination occurred leading to a loss of heterozygosity spanning the beta-tubulin locus. We quantified the heterozygosity of single nucleotide polymorphisms (SNPs) across the 7 contigs in our *N. parisii* assembly. For the four samples containing beta-tubulin homozygous mutations (S1-Alb, S2-Alb, S4-Alb, and S6-Alb), we observed a loss of heterozygosity on contig 3 from around the 100 kb position, until the end of the contig (Fig. 6). The beta-tubulin gene is located on contig 3 at position 0.45 Mb, so this loss of heterozygosity spanned the gene, and we did not observe a loss of heterozygosity around beta-tubulin for any of the other strains (Fig. 6). We also did not observe loss of heterozygosity for the albendazole-resistant strains containing homozygous beta-tubulin variants on any of the other contigs (Fig. 6). We did observe smaller regions of loss of heterozygosity on the right side of contig 1 for S3-Dex and part of the left side of contig 3 for S2-Dex and S3-Dex (Fig. 6). Together these results provide evidence for mitotic recombination events that resulted in homozygous beta-tubulin variants.

**Figure 6.**
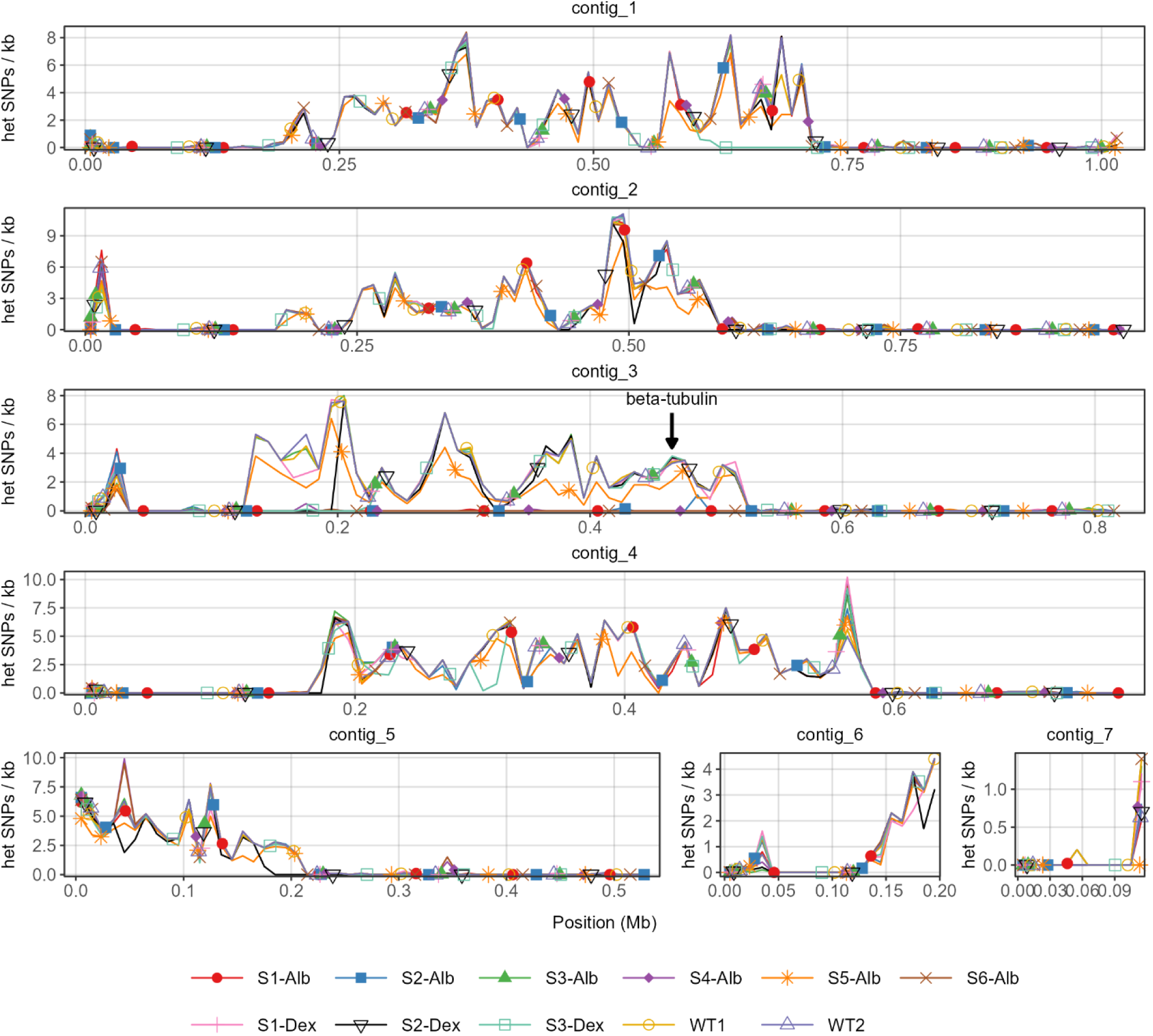
Loss of heterozygosity spanning the beta-tubulin locus occurs in albendazole-resistant *N. parisii* strains containing homozygous beta-tubulin variants. Heterozygous SNPs were determined for the nine drug-resistant *N. parisii* strains and two wild-type ERTm1 (WT1 and WT2) samples. The number of heterozygous SNPs per kb was calculated at 10 kb windows and mapped to the long-read *N. parisii* assembly. The location of the beta-tubulin locus on contig 3 is indicated with an arrow. The identity of the strains is indicated by the legend at the bottom.

## Discussion

To identify targets of microsporidia inhibitors, we developed a selection approach using *C. elegans* infected with *N. parisii*. We isolated six albendazole-resistant strains, all containing mutations in beta-tubulin, and two of these strains also contained mutations in alpha-tubulin. For dexrazoxane, we generated three resistant strains that all had mutations in topoisomerase II. Together our experiments demonstrate the ability to perform forward genetics in microsporidia, allowing for the association of genotype to phenotype. We also found evidence of mitotic recombination, observed through the loss of heterozygosity in strains with homozygous beta-tubulin mutations. Although all strains had mutations in genes known to confer resistance to the corresponding compound in other organisms, biochemical experiments with *N. parisii* beta-tubulin and topoisomerase II will be necessary to fully characterize the effects of these variants. Our data also suggest that in clinically relevant microsporidian species, the presence of homozygous and heterozygous mutations in beta-tubulin and alpha-tubulin should be monitored for potential signs of benzimidazole resistance.

Benzimidazoles have been demonstrated to inhibit beta-tubulin in many organisms and several positions have been described as providing resistance [24–28]. Position 198 is associated with benzimidazole resistance in parasitic nematodes and an E198K variant was shown to provide resistance in *C. elegans.* This variant was also detected in a benzimidazole-resistant mutant of the plant fungal pathogen *Botrytis cinerea* and the parasite *Giardia duodenalis* [24,25,28,29]. Two human-infecting microsporidia, *Enterocytozoon bieneusi* and *Vittaforma corneae,* have E198Q variants in beta-tubulin that are thought to be responsible for albendazole resistance in these species [6,7]. The Q134H variant is associated with benzimidazole resistance in the parasitic nematode *Ancylostoma caninum* and confers albendazole resistance when introduced into *C. elegans*. The Q134K variant was detected as a carbendazim resistance mutation in the wheat fungal pathogen *Zymoseptoria tritici* [30,31]. Although to the best of our knowledge, F20, A248, and A314 have not been associated with albendazole resistance, molecular docking suggests that F20 is part of the albendazole binding pocket and A248 and A314 are proposed to be in the binding site of the beta-tubulin inhibitor colchicine [32,33]. The variants we identified occur at evolutionarily conserved positions, with positions 20, 134, 198, and 314 conserved in *N. parisii*, *C. elegans*, yeast, and humans, and position 248 is conserved in all but yeast. At lower doses of benzimidazole, heterozygous beta-tubulin variants have been reported to show resistance in parasitic nematodes [27]. We observe that the homozygous variants are resistant to multiple benzimidazoles, unlike the heterozygous variants that are only resistant to albendazole, suggesting that performing selections with various benzimidazoles could result in different mutational spectra. We observe two albendazole-resistant strains that contain alpha-tubulin mutations. In *Saccharomyces cerevisiae,* several alanine scanning mutations in alpha-tubulin result in resistance to the benzimidazole analog benomyl and these mutations may indirectly provide drug resistance by increasing tubulin stability [26].

There are several mechanisms that could allow for the generation of homozygous variants in diploid organisms. Sexual recombination has been proposed in microsporidia as there are genomic signs of recombination as well as the presence of conserved meiotic genes, although sexual recombination has not been observed in laboratory settings [34–36]. Additionally, the asexual process of mitotic recombination can result in the loss of heterozygosity through gene conversion, mitotic crossovers, or break-induced replication [37]. In strains containing homozygous beta-tubulin mutations, we observe loss of heterozygosity only on the chromosome containing the beta-tubulin locus and not on other contigs, arguing that sexual recombination did not occur. Loss of heterozygosity over part of a chromosome that continues to the chromosomal terminus is associated with either a mitotic crossover or break-induced replication [37]. Mitotic recombination in yeast is both widespread and contributes to adaptive evolution of diploid organisms [38,39]. There are several examples of drug resistance occurring through homozygous variants generated through mitotic recombination in the pathogen fungi *Candida albicans* [40,41]. As heterozygous mutations in beta-tubulin have been shown to provide resistance to lower doses of benzimidazole, a likely scenario is that a *de novo* heterozygous mutation occurred, followed by mitotic recombination resulting in a homozygous variant which is resistant to high doses of albendazole [27].

Dexrazoxane has broad anti-microsporidia activity, being reported to inhibit six species [12,17,42]. The structure of yeast topoisomerase II in complex with dexrazoxane shows seven amino acids composing the binding pocket. Two of the mutations that occur in the dexrazoxane-resistant *N. parisii* strains, N142S and Y144H, are in this binding pocket [22]. Additionally, a mutation to serine at position Y165 (which corresponds to Y144 in yeast and *N. parisii*) in human TOP2A provides resistance to dexrazoxane [23]. All of the mutations we observed were heterozygous, consistent with previous findings that heterozygous mutations confer resistance to a dexrazoxane analog in Chinese hamster ovary cells [43]. To our knowledge, the third mutation we observed, V243M, has not been reported to be associated with resistance to dexrazoxane. Like the other two resistant variants, V243M is located in the ATPase domain and is conserved in human topoisomerase II. We observe several examples of short, internal losses of heterozygosity in the S2-Dex and S3-Dex strains. Dexrazoxane increases the rate of recombination in mammalian cells and stress in *C. albicans*, including exposure to antifungal drugs, increases the rate of mitotic recombination [44,45]. Previous experiments with another species of *Nematocida*, *N. ausubeli*, showed no evidence of loss of heterozygosity over multiple weeks of being passaged, suggesting that population-wide changes in heterozygosity do not occur due to adaptation to laboratory growth in *C. elegans* [35]. Together these observations suggest that treatment with dexrazoxane can cause an increase in loss of heterozygosity events in *N. parisii*. All the loss of heterozygosity events we observe on contig 3 likely involved recombination around the 100 kb position, suggesting a possible hotspot for recombination.

Our selection approach allows for both determining targets of novel inhibitors and which mutations in microsporidia arise for known targets. This approach is likely to also be useful for determining inhibitors of other stages of microsporidia infection such as those that block spore germination [12,15]. Beyond drug targets, this selection approach will be useful to determine the genetic basis for other phenotypes such as adaptation to host mutations that inhibit infection [46–48]. Additionally, there is a lack of verified selectable markers for drug resistance in microsporidia, and the dexrazoxane variants we describe are potential candidates for drug-selectable markers for microsporidia genomic modification [49].

## Acknowledgements

We thank Gio Dela Cruz, P. M. Shreenidhi, and Jonathan Tersigni for helpful comments on this manuscript. This work was supported by a Canadian Institutes of Health Research grant (no. 461807 to A. W. R.) and Q. H and X. M. were supported by awards from the China Scholarship Council.

## Competing interests

The authors declare that they have no competing interests.

## Data availability

Sequencing data is available under BioProject PRJNA1416473 and all experimental data is in Data S1.

## Author contributions

Q.H. and A. W. R. conceptualized the experiments. Q.H. performed all the drug selection and validation experiments. Q.H and A. W. R. analyzed the experimental data. A. W. R. performed genomic analysis of drug-resistant strains. X. M. generated sequencing data and assembled the *N. parisii* genome, with assistance from W. Z. The paper was predominantly written by A. W. R., with contributions from Q. H. and X. M. Mentorship was provided by G. P., J. C., and A. W. R. Funding was acquired by A. W. R. All authors contributed to paper editing.

## Materials and methods

### *C. elegans* maintenance

The wild-type *C. elegans* strain N2 was cultured on nematode growth media (NGM) plates with the *Escherichia coli* OP50 strain as a food source [50]. To obtain age-synchronized populations, worms were washed from plates and treated with a sodium hypochlorite/sodium hydroxide solution for bleaching. The released embryos were then incubated in M9 buffer at 21°C for 18–24 h to allow for hatching.

### Culturing of E. coli

*E. coli* OP50 was streaked from a frozen stock onto LB agar plates and incubated overnight at 37°C. A single colony was then used to inoculate a liquid LB culture, which was grown for 18 hours at 37°C with shaking. The resulting culture was concentrated 10-fold by centrifugation and stored at 4°C.

### *N. parisii* spore preparation

Stocks of *N. parisii* spores were generated using our liquid culture protocol [16]. Approximately 6,000 L1-stage *C. elegans* N2 worms were infected with *N. parisii* spores (10,000 spores/μL) in 3 mL of S-complete medium containing OP50 [51] and incubated with rotation at 21°C for 4 days in a 15 mL conical tube. The culture was then scaled up to 50 mL of S-complete medium with OP50 and 50,000 L1 worms, followed by 6 days of shaking incubation in a 250 mL flask. After harvesting, the infected worms were stored at-80°C. Spores were subsequently purified by mechanical disruption of the worms using 2 mm zirconia beads, and the homogenate was cleared by filtration through a 5 μm membrane (Millipore). The final *N. parisii* spore preparations, confirmed to be free of bacterial and fungal contamination, were stored at-80°C. Spore concentration was determined by counting DY96-stained spores with a Cell-VU sperm counting slide [14].

### Culturing of drug-resistant *N. parisii* spores

To select for drug resistance, age-synchronized L1 *C. elegans* (N2) were infected with wild-type *N. parisii* spores (10,000 spores/μL) in a culture medium containing a selected concentration of microsporidian inhibitors and incubated at 25°C for 6 days with rotation. The initial inhibitor concentration was determined by DY96 staining, requiring >50% of worms to exhibit moderate-to-severe infection. In subsequent cycles, a 1 mL aliquot from the previous culture was transferred to a fresh medium along with 10,000 L1-stage worms, and the inhibitor concentration was adjusted. The inhibitor concentration was increased if >50% of worms remained moderately-to-severely infected or decreased if >80% appeared uninfected. Cultures were maintained in 250 mL flasks at 25°C with shaking for 6 days until the OP50 food source was depleted. Strains were selected for resistance validation when >50% of worms were moderately-to-severely infected at inhibitor concentrations exceeding 60 μM.

### Validation of drug-resistant *N. parisii* strains

The resistance phenotype was subsequently validated using a standardized 24-well assay plate. In this assay, each well contained a total volume of 400 μL with 800 L1 worms and 10,000 spores/μL of the candidate *N. parisii* strain. Assays were performed in three biological replicates. Plates were sealed with breathable adhesive film, enclosed in parafilm-sealed boxes, and incubated at 21°C with shaking at 180 rpm for four days. Samples were then washed with M9/0.1% Tween 20, acetone-fixed, DY96-stained, and analyzed by fluorescence microscopy. A strain was confirmed as resistant if it maintained a moderate-to-severe infection level under these assay conditions.

### Elimination of bacterial and fungal contaminants during the culture of drug-resistant spores

To decontaminate drug-resistant *N. parisii* spore cultures, NGM plates were supplemented with a combination of antibiotics at the following final concentrations: chloramphenicol (100 μg/mL), carbenicillin (100 μg/mL), kanamycin (100 μg/mL), cefotaxime (100 μg/mL), gentamicin (10 μg/mL), and tetracycline (10 μg/mL) [52]. A small aliquot of the contaminated culture was applied to a corner of the antibiotic-containing plate and incubated overnight at 21°C. Any visible fungal hyphae or bacterial colonies, including the agar directly in contact with the inoculum, were aseptically excised. This process of incubation and excision was repeated periodically to remove regrowing contaminants. After 4 days, a clean section of the agar was transferred to a fresh antibiotic plate along with 200 L1-stage worms and incubated for 6 days at 21°C. If contamination persisted, the transfer process was repeated. Finally, a clean agar block was transferred to a standard NGM plate without antibiotics, supplemented with 200 L1 worms, and incubated for 4 days to confirm the absence of regrowth. Decontaminated cultures were harvested by suspending worms in ddH₂O for subsequent resistance screening.

### Extraction and whole-genome sequencing of *N. parisii* genomic DNA

*N. parisii*-infected worms, frozen at-80°C for at least 1 hour, were thawed and centrifuged at 1,400 rpm for 30 seconds. A 75 μL pellet was resuspended in 600 μL of pre-warmed (55°C) cell lysis buffer (10 mM Tris (pH 7.8), 5 mM EDTA, 0.5% SDS). Proteinase K (15-20 μL of 20 mg/mL) was added, and the mixture was incubated at 55°C for 1-2 hours with vortexing every 15 minutes until complete lysis was achieved. Following lysis, 2 μL of RNase A (100 mg/mL) was added, and the sample was incubated at 37°C for 45 minutes. After a 2-minute incubation on ice, the sample was centrifuged at 15,000 rpm for 3 minutes at 4°C to remove worm debris and bacteria. The supernatant was transferred to a new tube, mixed with 200 μL of 6M NaCl, and incubated on ice for 10 minutes. After centrifugation at 13,500 rpm for 12 minutes at 4°C, the supernatant was combined with 600 μL of isopropanol to precipitate the DNA. The sample was stored at-20°C for 24 hours, then centrifuged at 13,000 rpm for 12 minutes. The DNA pellet was washed three times with 600 μL of 70% ethanol, air-dried for 30 minutes, and dissolved in 30 μL of ultrapure water overnight at room temperature. Genomic DNA concentration was quantified using a Qubit BR system and stored at-20°C. DNA libraries were constructed by The Centre for Applied Genomics (TCAG) and sequenced using a NovaSeq X 10B flow cell to generate 150 bp paired-end reads.

### Targeted sequencing of potential drug resistance genes

Three primer pairs (0F: ATGAGGGAAATCGTACATTTAC; 350F: TGGATGTAATTAGAAAAGAGGC; 685F: GTTTCTGTTGTGATGAGTGG; R: TAGATTTATTAGATTTTAAATTACGG) were designed to amplify overlapping segments of the gene. PCR amplification was performed using the ERTm1 strain as a template in a 25 μL reaction mixture containing 1 μL of genomic DNA (250 ng/μL), 0.125 μL of Taq DNA Polymerase, 2 μL of primer mix, 0.5 μL of dNTPs, and 2.5 μL of 10× PCR Buffer. The amplification protocol consisted of an initial denaturation at 95°C for 3 min; 30 cycles of 95°C for 30 s, 50°C for 30 s, and 72°C for 2 min; followed by a final extension at 72°C for 5 min. PCR products were analyzed by electrophoresis on a 2% agarose gel and submitted for sequencing to TCAG.

### DNA extraction, long-read sequencing, and assembly of *N. parisii* strain ERTm1

Nematodes infected with microsporidia were collected by washing the plates with M9 solution, transferring the suspension to a 15 mL Falcon tube, and adjusting the volume to 14 mL with M9. The worms were washed three times with 14 mL of M9, centrifuged at 2,000 rpm for 2 minutes, and the supernatant discarded. After the final wash, the worm pellet was separated and stored at-80°C for at least 1 hour.

The frozen pellet was resuspended in 3 mL of Cell Lysis Solution (QIAGEN, Cat no./ID.158043) and incubated at 50°C for 1-2 hours, with gentle inversion every 30 minutes. Proteinase K (15 µL, QIAGEN, Cat no./ID.19131) and RNase A (15 µL, QIAGEN, Cat no./ID.158153) were added, followed by incubation at 37°C for 30 minutes. The lysate was then mixed with 1 mL of Protein Precipitation Solution, vortexed, and centrifuged. The supernatant was transferred to a new tube containing 3 mL of isopropanol, gently inverted, and centrifuged again. The resulting DNA pellet was washed with 70% ethanol, dried, and eluted overnight in 600 µL TE buffer at room temperature. DNA concentration was measured using a NanoDrop spectrophotometer.

To size select DNA, 3 µg of DNA was treated with a 2x Size Selection Solution (2.5% PVP360,000, 1.2 M NaCl, 20 mM Tris-HCl, pH 8) and centrifuged. DNA concentration was then quantified using Qubit High Sensitivity reagents (Invitrogen, Q32851) and a Qubit fluorometer. Size-selected DNA was subjected to DNA repair, end-prep, adapter ligation, and clean-up using the Ligation Sequencing Kit V14 (Oxford Nanopore, SQK-LSK114). The final library was eluted in 12 µL of Elution Buffer (EB).

The prepared library was sequenced using the MinION portable sequencing device (Oxford Nanopore) with R10.4.1 flow cells (FLO-MIN114), enabling direct DNA sequencing. Base calling was performed using the dna_r10.4.1_e8.2_400bps_sup@v5.0.0 model. Simplex and duplex base calling data were then extracted from the base called files using SAMtools (v1.13) [53]. Next, the extracted data were aligned to the host genomes (*C. elegans* and OP50) using minimap2 to remove host DNA and retain microsporidia DNA [54]. After alignment, reads with a Q score less than 15, shorter than 10,000 bp, and with the first and last 50 bp of each read were trimmed using NanoFilt (v2.6.0) to obtain high-quality filtered data. HERRO error correction was performed on the filtered data to correct sequencing errors, followed by genome assembly using Flye-2.9.4 to generate the *N. parisii* genome [55,56].

### Read mapping and variant calling

Paired-end sequencing reads were aligned to the long-read *N. parisii* genome assembly using BWA-MEM (v0.7.17) with default parameters [57]. Alignments were sorted and indexed using SAMtools (v1.13) [53]. Duplicate reads were marked using GATK (v4.6.2.0) MarkDuplicates [58]. Alignments were filtered and only primary alignments with mapping quality ≥30 were retained.

### Variant calling and annotation

Variants were called using GATK HaplotypeCaller (v4.6.2.0) in GVCF mode assuming diploid ploidy [58]. Per-sample GVCFs were combined using GATK CombineGVCFs and jointly genotyped using GenotypeGVCFs. Functional consequences were annotated using bcftools (v1.13) csq with gene models predicted by Prodigal (v2.6.3.0) [59,60]. Variants were filtered to require QUAL ≥30, QD ≥2.0, MQ ≥40, SOR ≤3.0, and strand bias (FS) ≤60 for SNPs or ≤200 for indels. Only variants containing missense mutations, frameshift mutations, or changes to stop codons were considered.

### Identification of variants associated with drug resistance

To identify candidate resistance mutations, we compared variant genotypes between either albendazole-resistant or dexrazoxane-resistant isolates and all other sequenced strains. Variants were classified as resistance-associated if they met the following criteria: (1) the variant was absent in the non-resistant samples (allele balance <1% at ≥20× coverage), and (2) the variant was present in at least one resistant isolate as either homozygous (≥98% allele balance, ≥50× coverage, ≥30 variant reads, genotype quality ≥50) or heterozygous (25–75% allele balance, ≥20 reads supporting each allele, ≥50× coverage, genotype quality ≥50). All variants are reported in Table S1.

### Read depth analysis

To assess copy number variation across the genome, read depth was calculated in 10 kb windows using mosdepth (v0.3.3) with a minimum mapping quality threshold of 30 [61]. For each sample, windowed depth values were normalized to the sample median to generate depth ratios, allowing comparison across samples with different sequencing depths.

### Loss of heterozygosity analysis

To detect regions of loss of heterozygosity (LOH), we analyzed the genome-wide distribution of heterozygous SNPs. We filtered SNPs to retain only biallelic sites using the same criteria described above. Heterozygous SNP density was calculated in 10 kb sliding windows across the genome and normalized to SNPs per kilobase.

## Supplementary materials

**Figure S1.**
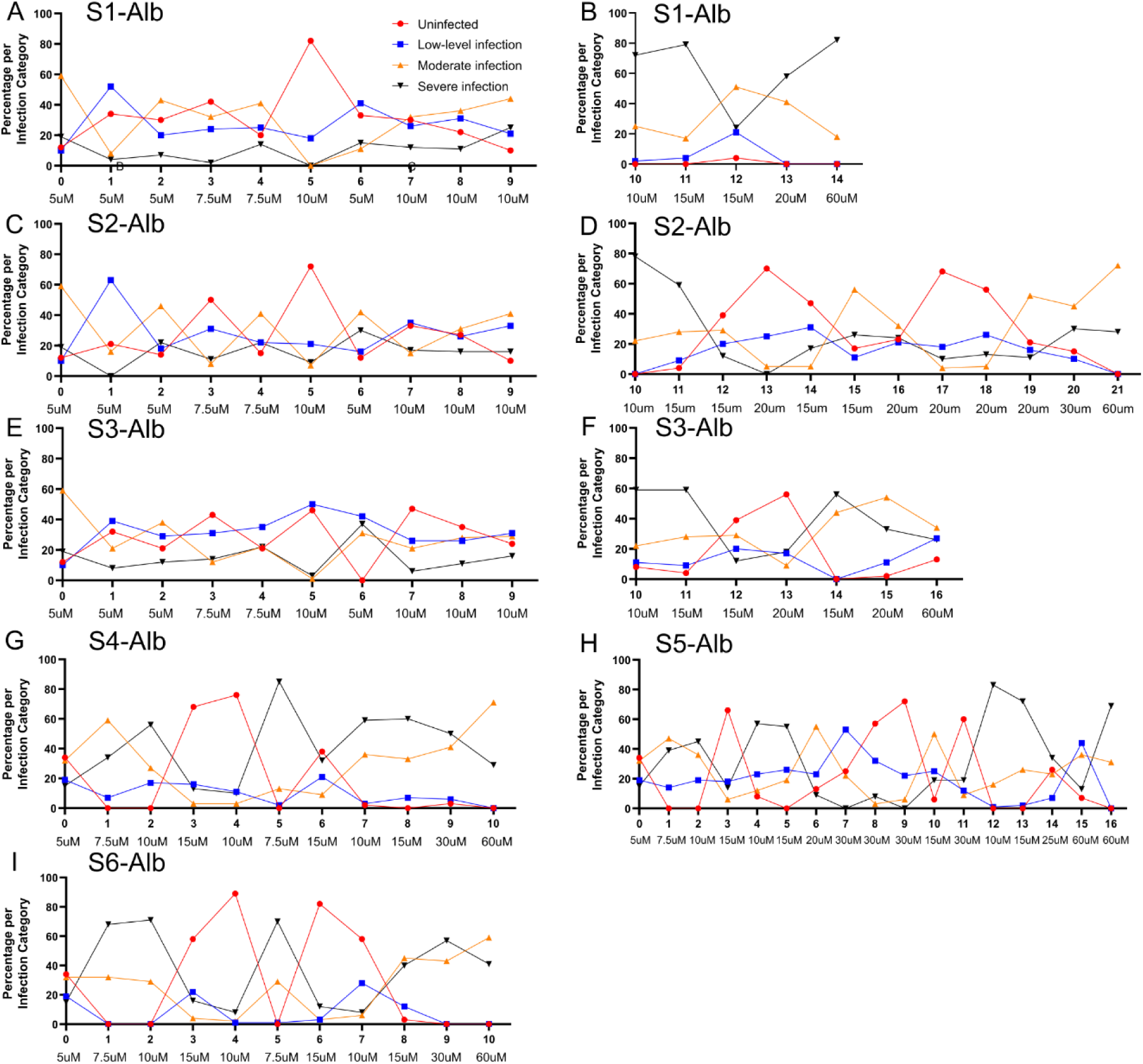
Relationship between drug concentration and infection severity in albendazole-resistant spore selections over multiple generations. A-I. For the initial (A, C, E; incubated at 21°C for 4 days) and optimized (B, D, F, G, H, I; incubated at 25°C for 6 days) protocols, a co-culture was established by adding 1 mL of infected nematode culture to 49 mL of 2× S-complete containing 10,000 L1 larvae. Infection status was assessed using DY96 and DAPI staining. The albendazole concentration was decreased if >80 % of adults were uninfected, increased if >50 % showed moderate to severe infection, and otherwise maintained. Cultures exhibiting >50 % moderate to severe infection at 60 µM albendazole were considered to harbor resistant spores and were subsequently validated by a 24-well continuous infection assay. Generation number is shown in bold under the x-axis along with the albendazole concentration.

**Figure S2.**
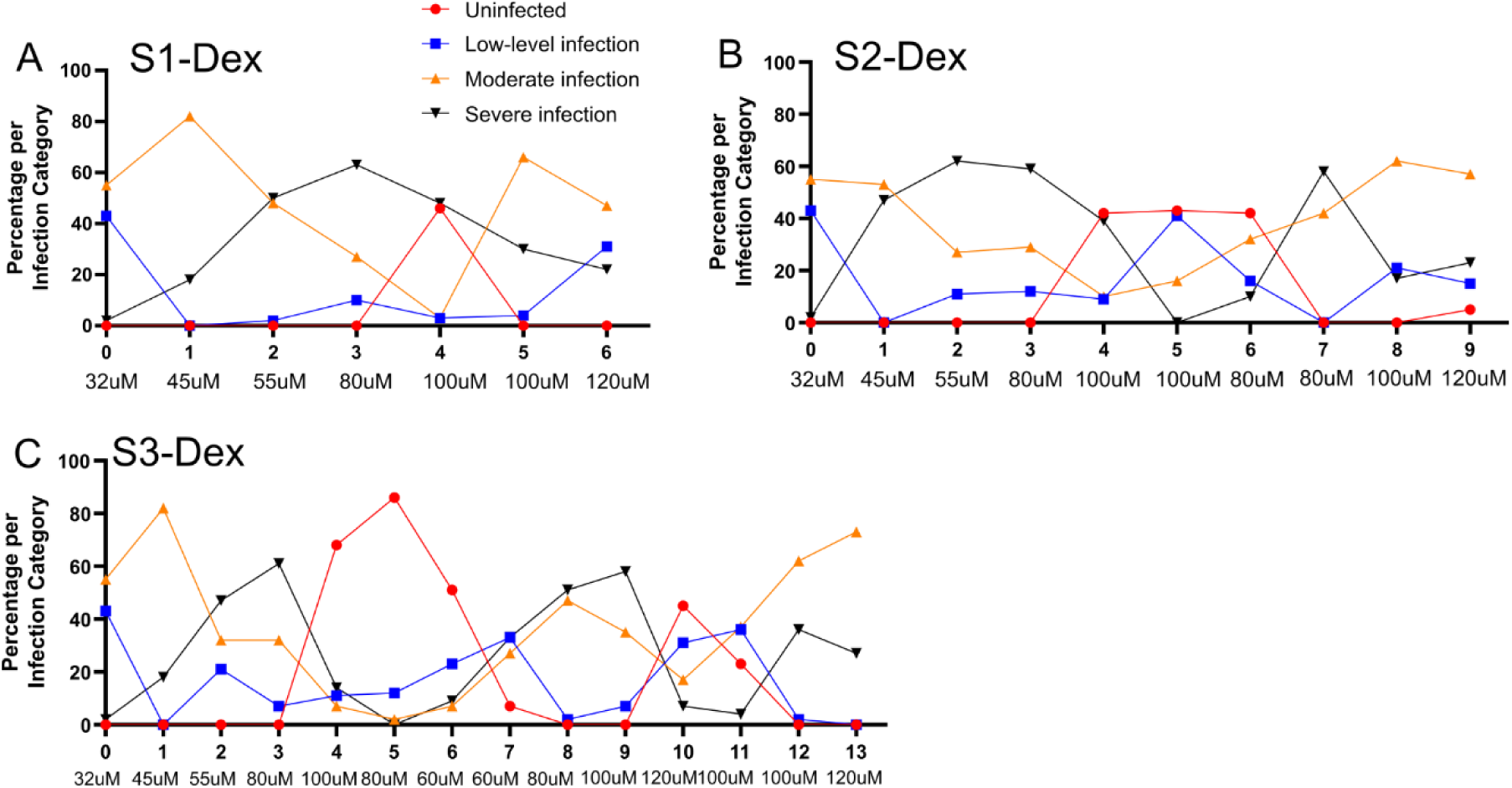
Relationship between drug concentration and infection severity in dexrazoxane-resistant spore selections over multiple generations. A-C. Application of the optimized resistant spore screening protocol. A co-culture was established by adding 1 mL of infected nematode culture to 49 mL of 2× S-complete containing 10,000 L1 worms, followed by incubation at 25 °C for 6 days. Infection status was assessed using DY96 and DAPI staining. The dexrazoxane concentration was decreased if >80% of adult worms were uninfected, increased if >50% exhibited moderate-to-severe infection, and maintained otherwise. Cultures showing >50% moderate-to-severe infection at 120 μM dexrazoxane were considered to harbor resistant spores and proceeded to validation by 24-well continuous infection assay. Generation number is shown in bold under the x-axis along with the dexrazoxane concentration.

**Figure S3.**
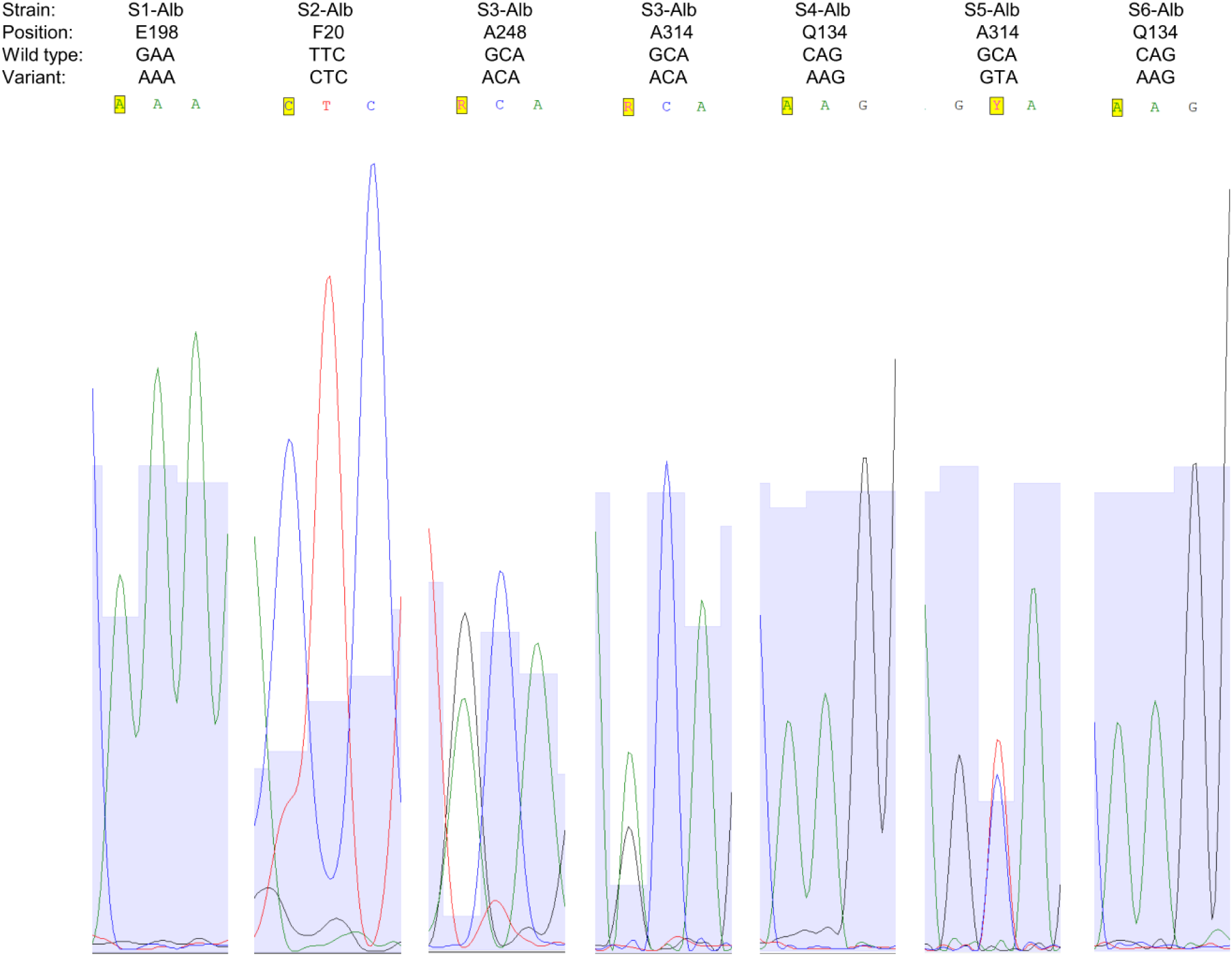
Sanger-sequencing validation of beta-tubulin mutations in albendazole-resistant *N. parisii* strains. The beta-tubulin gene from the albendazole-resistant strains was amplified using PCR and Sanger sequenced. The sequencing chromatograms for the codon containing the variants detected by whole genome sequencing are shown for each strain. The bases are colored with adenine in green, cytosine in blue, thymine in red, and guanine in black. The base changed in the resistant strains is highlighted in yellow. The position, the ERTm1 wild-type codon, and the variant observed in the resistant strain are shown above the chromatograms. APE was used to visualize chromatograms [65].

**Figure S4.**
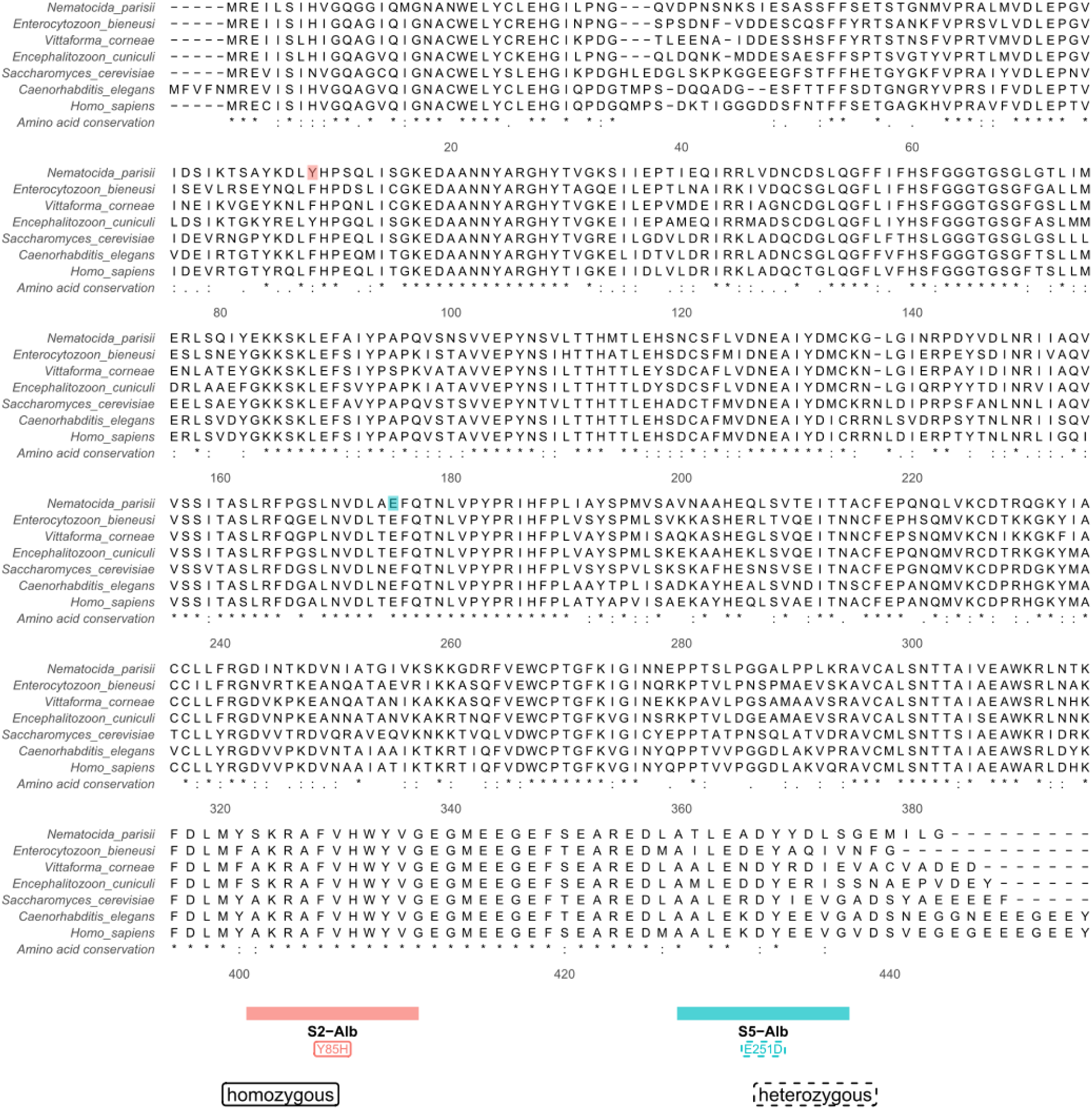
Two albendazole-resistant *N. parisii* strains contain either homozygous or heterozygous variants in alpha-tubulin. Multiple sequence alignment of alpha-tubulin sequences from microsporidia *(N. parisii*, *E. bieneusi*, *V. corneae*, and *E. cuniculi), S. cerevisiae*, *C. elegans*, and humans. Homozygous and heterozygous variants that were detected in two of the albendazole-resistant strains are annotated according to the legend at the bottom. Positions are numbered according to *S. cerevisiae* TUB1 and amino acid conservation is shown at the bottom of the alignment.

**Figure S5.**
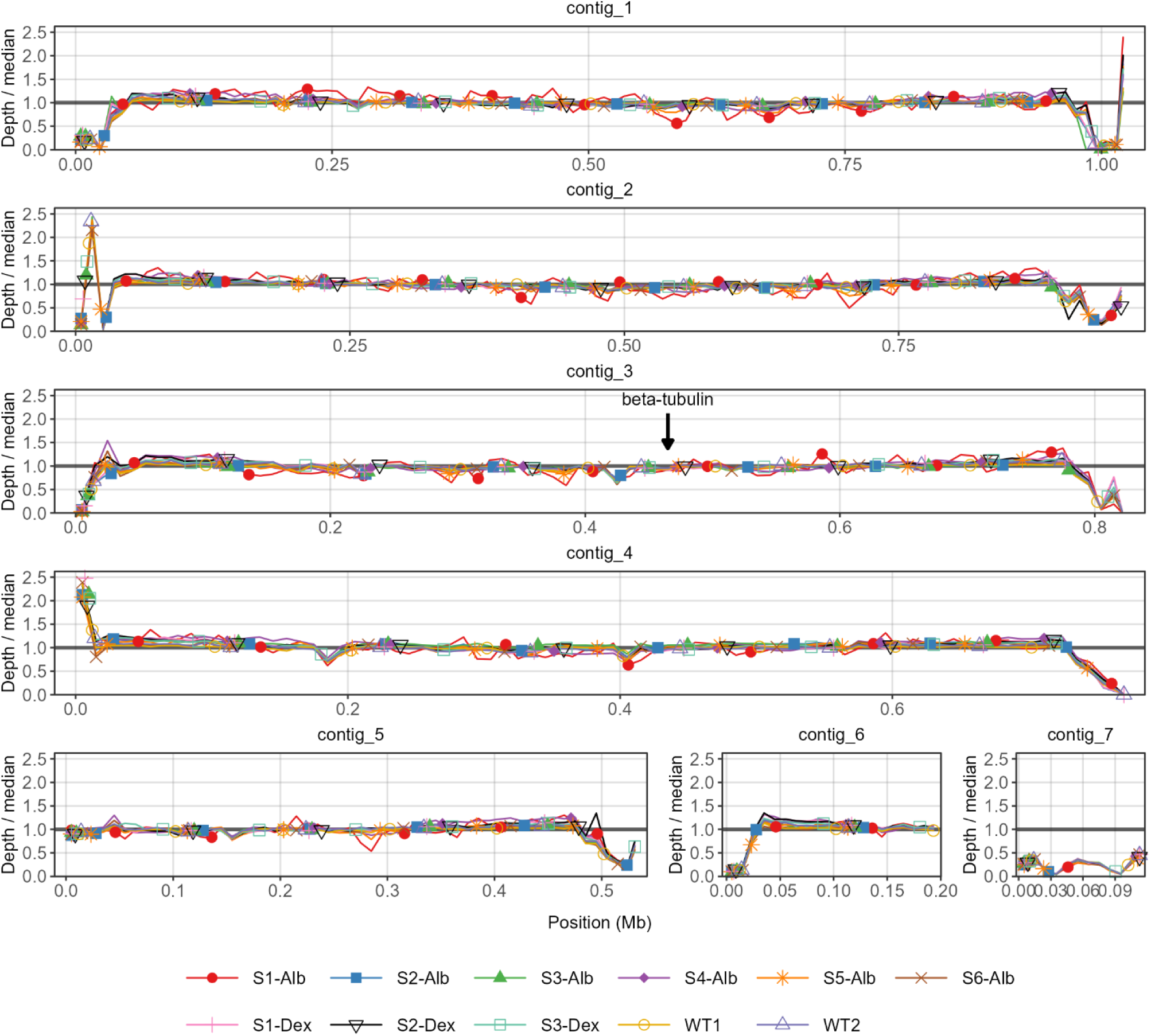
Read depth in drug-resistant *N. parisii* strains does not support aneuploidy. The read depth across the *N. parisii* genome assembly was determined for the nine drug-resistant *N. parisii* strains and two wild-type ERTm1 (WT1 and WT2) samples. The read depth at 10 kb windows divided by the median read depth for each sample at each contig is shown. The location of the beta-tubulin locus on contig 3 is indicated with an arrow. The identity of the strains is indicated by the legend at the bottom.

**Table S1. *N. parisii* homozygous and heterozygous variants with predicted protein-coding impacts that occur in albendazole-or dexrazoxane-resistant strains.**

**Data S1. All infection data from the study.**

## Notes

### Competing Interest Statement

The authors have declared no competing interest.

## References

1. Murareanu BM, Sukhdeo R, Qu R, Jiang J, Reinke AW. Generation of a Microsporidia Species Attribute Database and Analysis of the Extensive Ecological and Phenotypic Diversity of Microsporidia. mBio. 2021;12:. doi:10.1128/mbio.01490-21

2. Bojko J, Reinke AW, Stentiford GD, Williams B, Rogers MSJ, Bass D. Microsporidia: a new taxonomic, evolutionary, and ecological synthesis. Trends in Parasitology. 2022;38: 642–659. doi:10.1016/j.pt.2022.05.007

3. Han B, Pan G, Weiss LM. Microsporidiosis in Humans. Clinical Microbiology Reviews. 2021;34: e00010–20. doi:10.1128/CMR.00010-20

4. Franzen C. Microsporidia: A Review of 150 Years of Research. TOPARAJ. 2008;2: 1–34. doi:10.2174/1874421400802010001

5. Shinn A, Pratoomyot J, Griffiths D, Trọng T, Thanh Vu N, Jiravanichpaisal P, et al. Asian Shrimp Production and the Economic Costs of Disease. Asian Fisheries Science. 2018;31S. doi:10.33997/j.afs.2018.31.S1.003

6. Franzen C, Salzberger B. Analysis of the β-Tubulin Gene from Vittaforma corneae Suggests Benzimidazole Resistance. Antimicrobial Agents and Chemotherapy. 2008;52: 790–793. doi:10.1128/aac.00928-07

7. Akiyoshi DE, Weiss LM, Feng X, Williams B a. P, Keeling PJ, Zhang Q, et al. Analysis of the β-Tubulin Genes from Enterocytozoon bieneusi Isolates from a Human and Rhesus Macaque. Journal of Eukaryotic Microbiology. 2007;54: 38–41. doi:10.1111/j.1550-7408.2006.00140.x

8. Das BC, Chokkalingam P, Shareef MA, Shukla S, Das S, Saito M, et al. Methionine aminopeptidases: Potential therapeutic target for microsporidia and other microbes. Journal of Eukaryotic Microbiology. 2024;71: e13036. doi:10.1111/jeu.13036

9. Sak B, Brdíčková K, Holubová N, Květoňová D, Hlásková L, Kváč M. Encephalitozoon cuniculi Genotype III Evinces a Resistance to Albendazole Treatment in both Immunodeficient and Immunocompetent Mice. Antimicrobial Agents and Chemotherapy. 2020;64: doi:10.1128/aac.00058-20

10. Troemel ER, Felix M-A, Whiteman NK, Barriere A, Ausubel FM. Microsporidia are natural intracellular parasites of the nematode Caenorhabditis elegans. PLoS Biol. 2008;6: 2736–2752. doi:10.1371/journal.pbio.0060309

11. Gang SS, Lažetić V. Microsporidia: Pervasive natural pathogens of Caenorhabditis elegans and related nematodes. Journal of Eukaryotic Microbiology. 2024;71: e13027. doi:10.1111/jeu.13027

12. Murareanu BM, Antao NV, Zhao W, Dubuffet A, El Alaoui H, Knox J, et al. High-throughput small molecule screen identifies inhibitors of microsporidia invasion and proliferation in C. elegans. Nat Commun. 2022;13: 5653. doi:10.1038/s41467-022-33400-y

13. Balla KM, Luallen RJ, Bakowski MA, Troemel ER. Cell-to-cell spread of microsporidia causes Caenorhabditis elegans organs to form syncytia. Nature Microbiology. 2016;1: 16144.

14. Willis AR, Zhao W, Sukhdeo R, Wadi L, El Jarkass HT, Claycomb JM, et al. A parental transcriptional response to microsporidia infection induces inherited immunity in offspring. Sci Adv. 2021;7: eabf3114. doi:10.1126/sciadv.abf3114

15. Huang Q, Chen J, Pan G, Reinke AW. Screening of the Pandemic Response Box identifies anti-microsporidia compounds. PLOS Neglected Tropical Diseases. 2023;17: e0011806. doi:10.1371/journal.pntd.0011806

16. Huang Q, Brown LE, Pan G, Wei J, Porco JA, Chen J, et al. Identification of natural products and synthetic analogs which inhibit microsporidia spores and prevent infection. Journal of Invertebrate Pathology. 2026;214: 108492. doi:10.1016/j.jip.2025.108492

17. Huang Q, Jiang H, Wei J, Dou Y, Pan G, Chen J, et al. Small-molecule screen in C. elegans identifies benzenesulfonamides as inhibitors of microsporidia spores. npj Antimicrob Resist. 2025;3: 41. doi:10.1038/s44259-025-00116-0

18. Peirson M, Pernal SF. A Systematic Review of Fumagillin Field Trials for the Treatment of Nosema Disease in Honeybee Colonies. Insects. 2024;15: 29. doi:10.3390/insects15010029

19. Herman EH, Hasinoff BB, Steiner R, Lipshultz SE. A review of the preclinical development of dexrazoxane. Progress in Pediatric Cardiology. 2014;36: 33–38. doi:10.1016/j.ppedcard.2014.09.006

20. Hasinoff BB, Patel D, Wu X. The Role of Topoisomerase IIβ in the Mechanisms of Action of the Doxorubicin Cardioprotective Agent Dexrazoxane. Cardiovasc Toxicol. 2020;20: 312–320. doi:10.1007/s12012-019-09554-5

21. Balla KM, Andersen EC, Kruglyak L, Troemel ER. A wild C. elegans strain has enhanced epithelial immunity to a natural microsporidian parasite. PLoS Pathog. 2015;11: e1004583. doi:10.1371/journal.ppat.1004583

22. Classen S, Olland S, Berger JM. Structure of the topoisomerase II ATPase region and its mechanism of inhibition by the chemotherapeutic agent ICRF-187. Proceedings of the National Academy of Sciences. 2003;100: 10629–10634. doi:10.1073/pnas.1832879100

23. Wessel I, Jensen LH, Renodon-Corniere A, Sorensen TK, Nitiss JL, Jensen PB, et al. Human small cell lung cancer NYH cells resistant to the bisdioxopiperazine ICRF-187 exhibit a functional dominant Tyr165Ser mutation in the Walker A ATP binding site of topoisomerase IIα. FEBS Letters. 2002;520: 161–166. doi:10.1016/S0014-5793(02)02805-3

24. Dilks CM, Koury EJ, Buchanan CM, Andersen EC. Newly identified parasitic nematode beta-tubulin alleles confer resistance to benzimidazoles. International Journal for Parasitology: Drugs and Drug Resistance. 2021;17: 168–175. doi:10.1016/j.ijpddr.2021.09.006

25. Emery-Corbin SJ, Su Q, Tichkule S, Baker L, Lacey E, Jex AR. In vitro selection of Giardia duodenalis for Albendazole resistance identifies a β-tubulin mutation at amino acid E198K. Int J Parasitol Drugs Drug Resist. 2021;16: 162–173. doi:10.1016/j.ijpddr.2021.05.003

26. Richards KL, Anders KR, Nogales E, Schwartz K, Downing KH, Botstein D. Structure–Function Relationships in Yeast Tubulins. MBoC. 2000;11: 1887–1903. doi:10.1091/mbc.11.5.1887

27. Silvestre A, Cabaret J, Humbert J-F. Effect of benzimidazole under-dosing on the resistant allele frequency in Teladorsagia circumcincta (Nematoda). Parasitology. 2001;123: 103–111. doi:10.1017/S0031182001008009

28. Fan F, Li N, Li GQ, Luo CX. Occurrence of Fungicide Resistance in Botrytis cinerea from Greenhouse Tomato in Hubei Province, China. Plant Disease. 2016;100: 2414–2421. doi:10.1094/PDIS-03-16-0395-RE

29. Ghisi M, Kaminsky R, Mäser P. Phenotyping and genotyping of *Haemonchus contortus* isolates reveals a new putative candidate mutation for benzimidazole resistance in nematodes. Veterinary Parasitology. 2007;144: 313–320. doi:10.1016/j.vetpar.2006.10.003

30. Matsuzaki Y, Watanabe S, Harada T, Iwahashi F. Pyridachlometyl has a novel anti-tubulin mode of action which could be useful in anti-resistance management. Pest Management Science. 2020;76: 1393–1401. doi:10.1002/ps.5652

31. Venkatesan A, Castro PDJ, Morosetti A, Horvath H, Chen R, Redman E, et al. Molecular evidence of widespread benzimidazole drug resistance in Ancylostoma caninum from domestic dogs throughout the USA and discovery of a novel β-tubulin benzimidazole resistance mutation. PLOS Pathogens. 2023;19: e1011146. doi:10.1371/journal.ppat.1011146

32. Kenchappa R, Bodke YD, Telkar S, Aruna Sindhe M. Antifungal and anthelmintic activity of novel benzofuran derivatives containing thiazolo benzimidazole nucleus: an in vitro evaluation. J Chem Biol. 2017;10: 11–23. doi:10.1007/s12154-016-0160-x

33. Sahakyan H, Abelyan N, Arakelov V, Arakelov G, Nazaryan K. In silico study of colchicine resistance molecular mechanisms caused by tubulin structural polymorphism. PLOS ONE. 2019;14: e0221532. doi:10.1371/journal.pone.0221532

34. Lee SC, Corradi N, Byrnes EJ, Torres-Martinez S, Dietrich FS, Keeling PJ, et al. Microsporidia Evolved from Ancestral Sexual Fungi. Current Biology. 2008;18: 1675–1679. doi:10.1016/j.cub.2008.09.030

35. Cuomo CA, Desjardins CA, Bakowski MA, Goldberg J, Ma AT, Becnel JJ, et al. Microsporidian genome analysis reveals evolutionary strategies for obligate intracellular growth. Genome Res. 2012;22: 2478–2488. doi:10.1101/gr.142802.112

36. Khalaf A, Zhou C, Weber CC, Vancaester E, Sims Y, Makunin A, et al. Forty new genomes shed light on sexual reproduction and the origin of tetraploidy in Microsporidia. PLOS Biology. 2025;23: e3003446. doi:10.1371/journal.pbio.3003446

37. Symington LS, Rothstein R, Lisby M. Mechanisms and Regulation of Mitotic Recombination in Saccharomyces cerevisiae. Genetics. 2014;198: 795–835. doi:10.1534/genetics.114.166140

38. Magwene PM, Kayıkçı Ö, Granek JA, Reininga JM, Scholl Z, Murray D. Outcrossing, mitotic recombination, and life-history trade-offs shape genome evolution in Saccharomyces cerevisiae. Proceedings of the National Academy of Sciences. 2011;108: 1987–1992. doi:10.1073/pnas.1012544108

39. James TY, Michelotti LA, Glasco AD, Clemons RA, Powers RA, James ES, et al. Adaptation by Loss of Heterozygosity in Saccharomyces cerevisiae Clones Under Divergent Selection. Genetics. 2019;213: 665–683. doi:10.1534/genetics.119.302411

40. Coste A, Selmecki A, Forche A, Diogo D, Bougnoux M-E, d’Enfert C, et al. Genotypic Evolution of Azole Resistance Mechanisms in Sequential Candida albicans Isolates. Eukaryotic Cell. 2007;6: 1889–1904. doi:10.1128/ec.00151-07

41. Dunkel N, Blaß J, Rogers PD, Morschhäuser J. Mutations in the multi-drug resistance regulator MRR1, followed by loss of heterozygosity, are the main cause of MDR1 overexpression in fluconazole-resistant Candida albicans strains. Molecular Microbiology. 2008;69: 827–840. doi:10.1111/j.1365-2958.2008.06309.x

42. Zeferino TG, Silva LM. Dexrazoxane as a viable microsporidia control agent in *Anopheles gambiae*. Acta Tropica. 2025;266: 107633. doi:10.1016/j.actatropica.2025.107633

43. Sehested M, Wessel I, Jensen LH, Holm B, Oliveri RS, Kenwrick S, et al. Chinese Hamster Ovary Cells Resistant to the Topoisomerase II Catalytic Inhibitor ICRF-159: A Tyr49Phe Mutation Confers High-Level Resistance to Bisdioxopiperazines1. Cancer Res. 1998;58: 1460–1468.

44. Forche A, Abbey D, Pisithkul T, Weinzierl MA, Ringstrom T, Bruck D, et al. Stress Alters Rates and Types of Loss of Heterozygosity in Candida albicans. mBio. 2011;2: doi:10.1128/mbio.00129-11

45. Jensen LH, Dejligbjerg M, Hansen LT, Grauslund M, Jensen PB, Sehested M. Characterisation of cytotoxicity and DNA damage induced by the topoisomerase II-directed bisdioxopiperazine anti-cancer agent ICRF-187 (dexrazoxane) in yeast and mammalian cells. BMC Pharmacol. 2004;4: 31. doi:10.1186/1471-2210-4-31

46. Tamim El Jarkass H, Mok C, Schertzberg MR, Fraser AG, Troemel ER, Reinke AW. An intestinally secreted host factor promotes microsporidia invasion of C. elegans. Lourido S, editor. eLife. 2022;11: e72458. doi:10.7554/eLife.72458

47. Jarkass HTE, Castelblanco S, Kaur M, Wan YC, Ellis AE, Sheldon RD, et al. The Caenorhabditis elegans bacterial microbiome influences microsporidia infection through nutrient limitation and inhibiting parasite invasion. bioRxiv; 2024. p. 2024.06.05.597580. doi:10.1101/2024.06.05.597580

48. Reddy KC, Dror T, Sowa JN, Panek J, Chen K, Lim ES, et al. An Intracellular Pathogen Response Pathway Promotes Proteostasis in C. elegans. Curr Biol. 2017;27: 3544–3553.e5. doi:10.1016/j.cub.2017.10.009

49. Reinke AW, Troemel ER. The Development of Genetic Modification Techniques in Intracellular Parasites and Potential Applications to Microsporidia. PLoS Pathog. 2015;11: e1005283. doi:10.1371/journal.ppat.1005283

50. Lewis JA, Fleming JT. Basic culture methods. Methods in cell biology. 1995;48: 3–29.

51. Hibshman JD, Webster AK, Baugh LR. Liquid-culture protocols for synchronous starvation, growth, dauer formation, and dietary restriction of *Caenorhabditis elegans*. STAR Protocols. 2021;2: 100276. doi:10.1016/j.xpro.2020.100276

52. Wadi L, Jarkass HTE, Tran TD, Islah N, Luallen RJ, Reinke AW. Genomic and phenotypic evolution of nematode-infecting microsporidia. PLOS Pathogens. 2023;19: e1011510. doi:10.1371/journal.ppat.1011510

53. Li H, Handsaker B, Wysoker A, Fennell T, Ruan J, Homer N, et al. The Sequence Alignment/Map format and SAMtools. Bioinformatics. 2009;25: 2078–2079. doi:10.1093/bioinformatics/btp352

54. Li H. Minimap2: pairwise alignment for nucleotide sequences. Bioinformatics. 2018;34: 3094–3100. doi:10.1093/bioinformatics/bty191

55. Stanojević D, Lin D, Nurk S, Sessions PF de, Šikić M. Telomere-to-Telomere Phased Genome Assembly Using HERRO-Corrected Simplex Nanopore Reads. bioRxiv; 2024. p. 2024.05.18.594796. doi:10.1101/2024.05.18.594796

56. Kolmogorov M, Yuan J, Lin Y, Pevzner PA. Assembly of long, error-prone reads using repeat graphs. Nat Biotechnol. 2019;37: 540–546. doi:10.1038/s41587-019-0072-8

57. Li H, Durbin R. Fast and accurate short read alignment with Burrows–Wheeler transform. Bioinformatics. 2009;25: 1754–1760. doi:10.1093/bioinformatics/btp324

58. McKenna A, Hanna M, Banks E, Sivachenko A, Cibulskis K, Kernytsky A, et al. The Genome Analysis Toolkit: A MapReduce framework for analyzing next-generation DNA sequencing data. Genome Res. 2010;20: 1297–1303. doi:10.1101/gr.107524.110

59. Hyatt D, Chen G-L, LoCascio PF, Land ML, Larimer FW, Hauser LJ. Prodigal: prokaryotic gene recognition and translation initiation site identification. BMC Bioinformatics. 2010;11: 119. doi:10.1186/1471-2105-11-119

60. Danecek P, Bonfield JK, Liddle J, Marshall J, Ohan V, Pollard MO, et al. Twelve years of SAMtools and BCFtools. Gigascience. 2021;10: giab008. doi:10.1093/gigascience/giab008

61. Pedersen BS, Quinlan AR. Mosdepth: quick coverage calculation for genomes and exomes. Bioinformatics. 2018;34: 867–868. doi:10.1093/bioinformatics/btx699

62. Wessel I, Jensen LH, Jensen PB, Falck J, Rose A, Roerth M, et al. Human Small Cell Lung Cancer NYH Cells Selected for Resistance to the Bisdioxopiperazine Topoisomerase II Catalytic Inhibitor ICRF-187 Demonstrate a Functional R162Q Mutation in the Walker A Consensus ATP Binding Domain of the α Isoform1. Cancer Res. 1999;59: 3442–3450.

63. Patel S, Jazrawi E, Creighton AM, Austin CA, Fisher LM. Probing the Interaction of the Cytotoxic Bisdioxopiperazine ICRF-193 with the Closed Enzyme Clamp of Human Topoisomerase IIα. Molecular Pharmacology. 2000;58: 560–568. doi:10.1016/S0026-895X(24)12421-2

64. Jensen LH, Wessel I, Møller M, Nitiss JL, Sehested M, Jensen PB. N-terminal and core-domain random mutations in human topoisomerase II α conferring bisdioxopiperazine resistance. FEBS Letters. 2000;480: 201–207. doi:10.1016/S0014-5793(00)01934-7

65. Davis MW, Jorgensen EM. ApE, A Plasmid Editor: A Freely Available DNA Manipulation and Visualization Program. Frontiers in Bioinformatics. 2022;2. Available: https://www.frontiersin.org/article/10.3389/fbinf.2022.818619

